# Processing of eukaryotic Okazaki fragments by redundant nucleases can be uncoupled from ongoing DNA replication *in vivo*

**DOI:** 10.1101/384503

**Authors:** Malik Kahli, Joseph S. Osmundson, Rani Yeung, Duncan J. Smith

## Abstract

Prior to ligation, each Okazaki fragment synthesized on the lagging strand in eukaryotes must be nucleolytically processed. Nuclease cleavage takes place in the context of 5’ flap structures generated via strand-displacement synthesis by DNA polymerase delta. At least three DNA nucleases: Rad27 (Fen1), Dna2, and Exo1, have been implicated in processing Okazaki fragment flaps. However, neither the contributions of individual nucleases to lagging-strand synthesis nor the structure of the DNA intermediates formed in their absence have been clearly defined *in vivo.* By conditionally depleting lagging-strand nucleases and directly analyzing Okazaki fragments synthesized *in vivo* in *S. cerevisiae*, we conduct a systematic evaluation of the impact of Rad27, Dna2 and Exo1 on lagging-strand synthesis. We find that Rad27 processes the majority of lagging-strand flaps, with a significant additional contribution from Exo1 but not from Dna2. When nuclease cleavage is impaired, we observe a reduction in strand-displacement synthesis as opposed to the widespread generation of long Okazaki fragment 5’ flaps, as predicted by some models. Further, using cell cycle-restricted constructs, we demonstrate that both the nucleolytic processing and the ligation of Okazaki fragments can be uncoupled from DNA replication and delayed until after synthesis of the majority of the genome is complete.

## INTRODUCTION

The synthesis of each Okazaki fragment requires several distinct enzymatic activities (Burgers, 2009). The primase component of the Pol α-primase complex synthesizes a short RNA primer, which is further extended by the error-prone DNA polymerase α (Pol α) (Nethanel and Kaufmann, 1990; Singh et al., 1986). The high-fidelity,PCNA-associated polymerase δ (Pol δ) is subsequently loaded onto the 3’ terminus of the initiating fragment in a reaction catalyzed by the RFC clamp loader (Maga et al., 2000). Pol δ synthesizes DNA to the 5’ end of the preceding fragment and continues beyond this point (Garg et al., 2004), generating a 5’ flap structure. The displaced 5’ flap serves as a substrate for nucleases (Kao et al., 2004), which cleave to generate a nick between Okazaki fragment termini. Iterative rounds of extension, followed by flap cleavage or nick regeneration by the 3’-5’ exonuclease activity of Pol δ (Garg et al., 2004), maintain a ligatable nick that persists until it is sealed by DNA ligase I, encoded by the *CDC9* gene in *S. cerevisiae* (Johnston and Nasmyth, 1978). Nuclease cleavage during Okazaki fragment biogenesis represents an extremely abundant DNA transaction – one that must occur tens of thousands of times during each S-phase in *S. cerevisiae* and millions of times per human cell division.

The nucleases Rad27 and Dna2 have been proposed to cleave the majority of flaps during Okazaki fragment maturation. Genetic and biochemical work has given rise to a ‘two-nuclease’ model (Balakrishnan et al., 2010; Burgers, 2009). According to this model, RNase H2 first removes most of the RNA primer (Qiu et al., 1999). Subsequently, iterative extension by Pol δ is followed by immediate cleavage of short DNA flaps by Rad27. If Pol δ extension outpaces Rad27 cleavage, Dna2 is required to process the resulting long flap. Rad27 and Dna2 show distinct substrate requirements *in vitro*. Rad27 readily cleaves short flaps; longer flaps are competent to bind RPA and thereby become refractory to Rad27 cleavage (Rossi and Bambara, 2006). Long, RPA-coated flaps are optimal substrates for Dna2 *in vitro.* Dna2 cleaves to leave a short flap such that RPA dissociates and Rad27 can cleave (Ayyagari et al., 2003; Gloor et al., 2012). However, recent reports indicate that Dna2 activity is sufficient to process 5’ flaps into ligatable nicks *in vitro* (Levikova and Cejka, 2015). Genetic data suggest additional redundancy in lagging-strand processing, and point to the likely involvement of Exo1 as a third Okazaki nuclease. *In vivo* in *S. cerevisiae*, neither *RAD27* nor *DNA2* is strictly essential for replication or viability (Budd et al., 2011), and the temperature-sensitive phenotypes of both *rad27Δ* and *dna2-1* can be suppressed by overexpression of *EXO1* (Budd and Campbell, 2000; Budd et al.,2005; Parenteau and Wellinger, 1999). Despite the apparent contribution of at least three deoxyribonucleases to lagging-strand processing, the normal contribution of each nuclease has not been clearly defined *in vivo*.

It is currently unclear whether impaired lagging-strand processing *in vivo* would predominantly give rise to reduced strand displacement by Pol δ, or to residual Okazaki fragment 5’ flaps. *In vitro* data suggests that strand-displacement synthesis should be dramatically reduced by a failure to cleave Okazaki fragment termini; instead, idling by Pol δ should maintain genomic nicks after limited strand displacement (Garg et al., 2004; Stodola and Burgers, 2016). However, electron microscopy analysis of DNA purified from *S. pombe* with lagging-strand processing defects detected the widespread accumulation of long flap structures (Liu et al., 2017). Furthermore, the ability of the replisome to bypass damage *in vivo* (Callegari et al., 2010; Lopes et al., 2006) and *in vitro* (Taylor and Yeeles, 2018) suggests that DNA repair can occur after bulk DNA synthesis. This model is supported by additional *in vivo* evidence – for example the ability of *S. cerevisiae* to defer post-replication repair to G2/M (Daigaku et al., 2010). Lagging-strand synthesis must inevitably occur at the same time as the leading strand, but the extent to which Okazaki fragment processing must be coupled to ongoing replication has not been determined.

Here, we conduct a systematic analysis of Okazaki fragment synthesis and processing in *S. cerevisiae* while depleting the lagging-strand nucleases Rad27, Dna2 and Exo1 in all possible combinations. Our data are consistent with a model whereby the extent of strand-displacement by Pol δ is severely reduced in the absence of nuclease cleavage. Rad27 cleaves most lagging-strand flaps, Exo1 serves as a redundant processing factor when Rad27 is absent, and the contribution of Dna2 to lagging-strand processing in the uniquely mappable regions of the *S. cerevisiae* genome is limited. Further, we show that cells remain viable when Okazaki fragment ligation is deferred until after bulk DNA synthesis in each cell cycle; nucleolytic processing of the lagging strand can be similarly deferred, but only if the accumulation of single-stranded DNA is mitigated during S-phase.

## RESULTS

### Systematic depletion of Okazaki nucleases and analysis of lagging-strand products

*DNA2* is an essential gene in *S. cerevisiae*, and *rad27Δ* mutants have several undesirable phenotypes including slow growth and elevated mutation rate (Reagan et al., 1995). In addition, genetic interactions between lagging-strand nucleases render double- and triple-mutant strains inviable (Budd et al., 2005). Therefore, we used the anchor away conditional nuclear depletion strategy to analyze Okazaki fragment synthesis in the absence of Rad27 and Dna2. Anchor away depletion is mediated by dimerization of FRB-tagged nuclear proteins with FKBP-tagged ribosomal subunits, and rapidly depletes proteins from the nucleus (Haruki et al., 2008). We constructed a panel of *S. cerevisiae* strains carrying all possible combinations of *exo1Δ*, *RAD27-FRB* and *DNA2-FRB*, in combination with an FRB-tagged allele of *CDC9* for simultaneous co-depletion of DNA ligase. All proteins were tagged at the C-terminus. Two biological replicates of each strain were obtained via sporulation of a diploid strain heterozygous at the *EXO1*, *DNA2* and *RAD27* loci and homozygous at the *CDC9* locus. Conditional nuclear depletion of Rad27 or Dna2 recapitulated the reported phenotype of the null mutants (Fig. 1A). *DNA2-FRB* strain were inviable upon rapamycin treatment, and *RAD27-FRB* strains were inviable at 37˚C in the presence of rapamycin. Double (Fig. 1A) and triple (not shown) nuclease mutants were inviable upon rapamycin treatment as expected.

**Figure 1.**
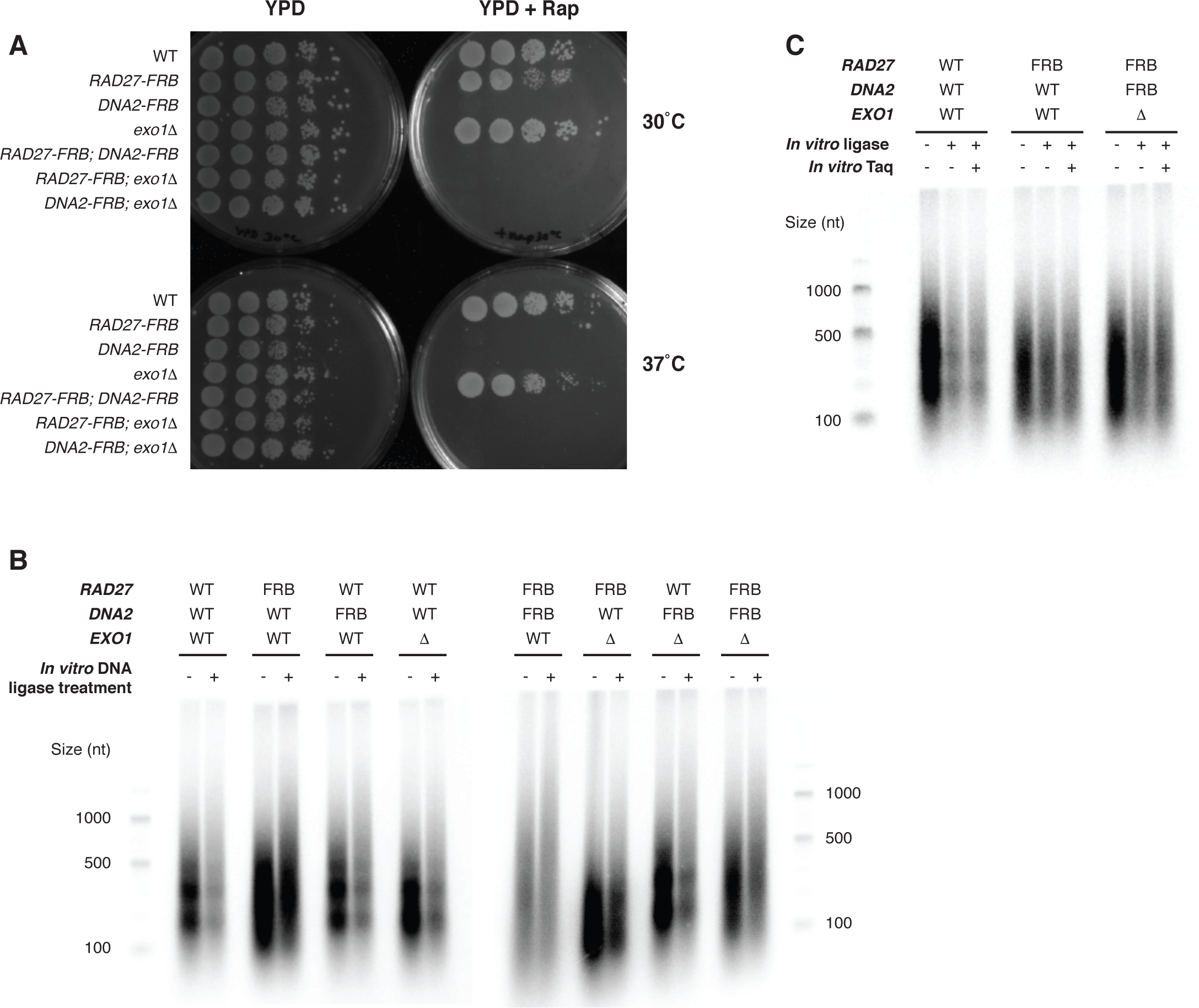
Enrichment of ligation-competent Okazaki fragments from strains lacking both lagging-strand nucleases and DNA ligase I. **A.** Spot tests of indicated strains on YPD ± 2 υg/ml rapamycin at 30˚C or 37˚C, as indicated, for single and double mutant strains analyzed in this study. Addition of rapamycin depletes FRB-tagged proteins from the nucleus. **B.** End-labeled Okazaki fragments obtained after 1h rapamycin treatment to deplete DNA ligase I and FRB-tagged nucleases. Where indicated, DNA was treated with T4 DNA ligase after purification but before labeling to assess the proportion of Okazaki fragments poised for ligation. WT denotes an otherwise wild-type *CDC9-FRB* strain. **C.** End-labeled Okazaki fragments, as in (B), for WT, *RAD27-FRB* and *RAD27-FRB DNA2-FRB*; *exo1*Δ strains. DNA was treated or mock-treated with Taq polymerase and DNA ligase after purification, as indicated.

Okazaki fragments in *S. cerevisiae* are sized according to the nucleosome repeat length due to interactions between Pol δ and nascent chromatin behind the replication fork, and are poised for ligation after purification (Smith and Whitehouse, 2012). We analyzed the size distribution of fragments from asynchronous cultures of strains depleted for DNA ligase and either one, two, or all three nucleases, and further analyzed the extent to which these fragments were competent for ligation after purification of genomic DNA (Fig. 1B, (Smith and Whitehouse, 2012)).

Okazaki fragment synthesis in the absence of one, two or three of the Okazaki nucleases did not lead to a gross change in fragment length (Fig. 1B). However, depletion of Rad27 from the nucleus consistently led to a loss of the nucleosome patterning observed in Rad27-proficient strains. In all strains, even in the absence of Rad27, Dna2 and Exo1, treatment with T4 DNA ligase after DNA purification resulted in a loss of Okazaki fragment end-labeling (Fig. 1B).

Therefore, the majority of Okazaki fragments in all strains are poised for ligation. The generation of Okazaki fragments bounded by ligatable nicks in the absence of processing nucleases is most consistent with a loss of strand-displacement by Pol δ. The alternative scenario, in which Pol δ carries out extensive strand-displacement in the absence of nuclease cleavage, would result in a large number of 5’ flap structures. To test whether such flap structures might be present, we treated purified DNA from wild-type, *RAD27-FRB* and *RAD27-FRB; DNA2-FRB; exo1Δ* strains with Taq polymerase prior to *in vitro* ligation (Fig. 1C). Taq polymerase has flap endonuclease activity (Lyamichev et al., 1993), and cleavage of 5’ flaps would therefore increase the proportion of Okazaki termini that can be processed by T4 DNA ligase. No increase in ligatability was observed upon Taq treatment, consistent with reduced strand-displacement as opposed to widespread 5’ flap formation when Okazaki fragment processing is perturbed *in vivo*.

### Okazaki fragment processing of nucleosomal DNA *in vivo*

In wild-type *S. cerevisiae*, Okazaki fragment termini are normally enriched at nucleosome dyads; a shift in this distribution towards the replication-fork-proximal edge of the nucleosome indicates reduced strand-displacement (Osmundson et al., 2017; Smith and Whitehouse, 2012). To directly investigate the extent of strand-displacement synthesis by Pol δ and nucleolytic Okazaki fragment processing in nucleosomal DNA, we purified and sequenced Okazaki fragments from asynchronous cultures (Smith and Whitehouse, 2012) after co-depletion of Cdc9 and all combinations of nucleases for 1h. Distributions of Okazaki fragment termini around nucleosome dyads are shown in Figure 2. All data were highly reproducible across two biological replicates. Data from the second replicate corresponding to Figures 2 & 3 are shown in Figure S1. For each sample, data were normalized to the maximum signal in the range to facilitate comparison between samples.

**Figure 2.**
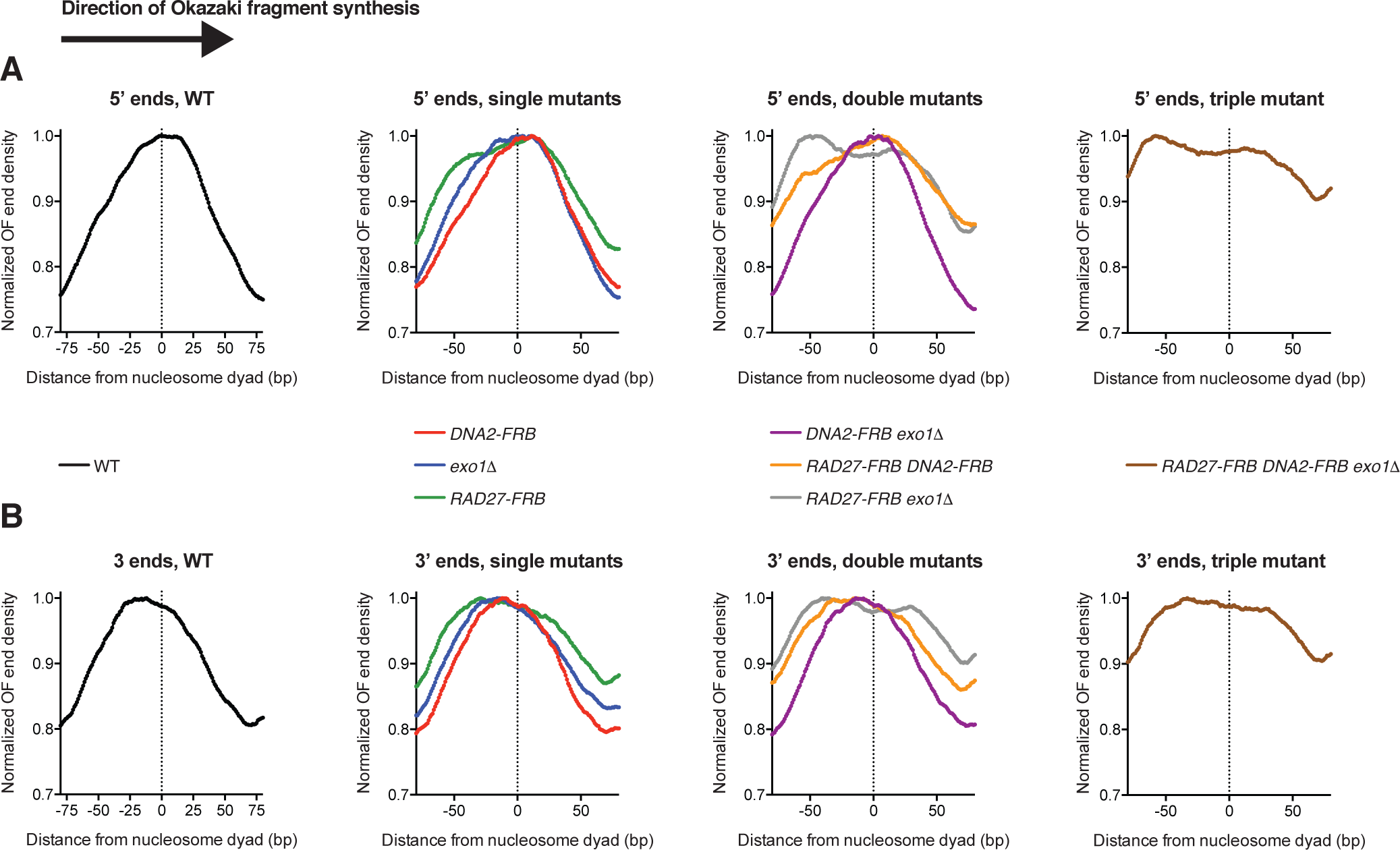
Contributions of individual nucleases to Okazaki fragment processing in the context of nucleosomal DNA. **A.** Distribution of Okazaki fragment 5’ termini around consensus dyad locations of the top 50% most highly occupied nucleosomes in the S. cerevisiae genome (Jiang and Pugh, 2009). WT, single mutants, double mutants and the triple mutant are displayed in separate graphs for clarity. Data are presented such that Okazaki fragment synthesis proceeds from left to right. Data are normalized to the maximum signal in the range, and smoothed to 5 bp. **B.** Distribution of Okazaki fragment 3’ termini around nucleosome dyads, as in (A).

Consistent with the observation that Okazaki fragments are competent for ligation even when processing nucleases are depleted, fragment 5’ (Fig. 2A) and 3’ (Fig. 2B) termini showed similar distributions within each sample. If strand displacement were occurring in the absence of nuclease cleavage, 3’ ends would show a wild-type distribution with respect to nucleosomes, while 5’ ends would shift to more upstream locations. Depletion of Rad27 led to a pronounced shift in both 5’ and 3’ termini towards the replication-fork proximal edge of the nucleosome. Deletion of *EXO1* produced a lesser shift in the same direction, while depletion of Dna2 did not significantly impact Okazaki fragment terminus location. Additional depletion of Dna2 did not dramatically alter the location of termini in *RAD27-FRB* or *exo1Δ* strains. However, cells lacking both Exo1 and Rad27 activity showed an additional shift compared to each single mutant, consistent with redundancy between the two nucleases. Okazaki fragment distributions in the *RAD27-FRB DNA2-FRB exo1Δ* strain were similar to those in *RAD27-FRB exo1Δ* (Fig. 2A&B). Taken together, this suggests that depletion of Dna2 to levels that do not support viability does not have a significant impact on nucleosomal Okazaki fragment processing throughout most of the genome, even when one or both of the other Okazaki nucleases are absent. The distribution of Okazaki fragment termini supports a model whereby Rad27 processes the majority of 5’ flaps, with a minor contribution from Exo1 and an extremely limited role for Dna2.

### Okazaki fragment processing of non-nucleosomal DNA *in vivo*

In nucleosome-free regions, Okazaki fragment termini are normally enriched immediately upstream of binding sites for the transcription factors Abf1, Reb1 and Rap1. These transcription factors re-associate quickly following DNA replication, and appear to represent ‘hard’ barriers to Pol δ in contrast to the ‘soft’ barrier activity of nucleosomes (Smith and Whitehouse, 2012). We analyzed the distribution of Okazaki fragment termini around these transcription factor binding sites, as a means to investigate strand-displacement and processing on non-nucleosomal templates (Fig. 3). Data were normalized to the median signal within the range in order to assess the propensity of Pol δ to terminate specifically at transcription-factor binding sites as opposed to nearby sequences.

**Figure 3.**
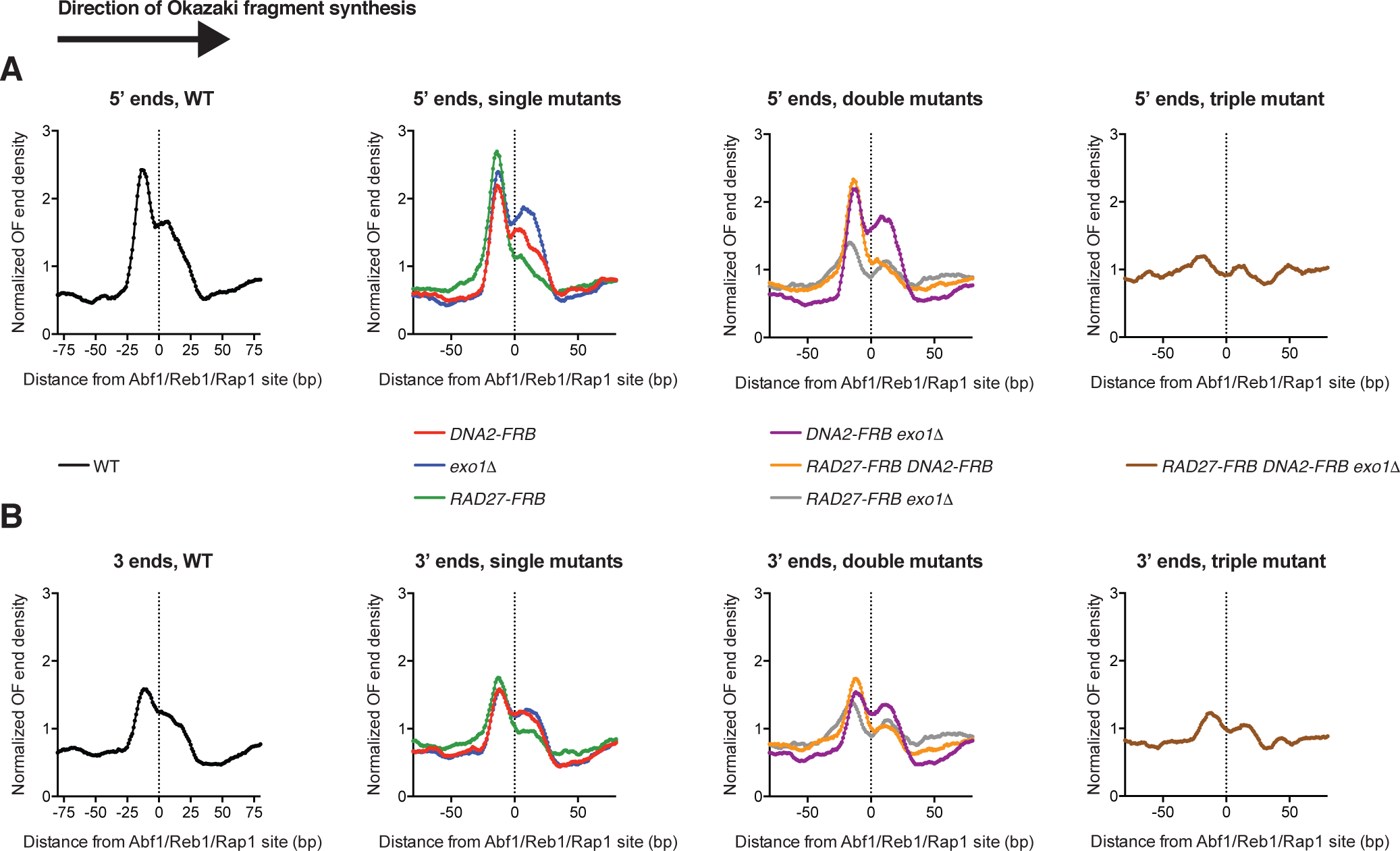
Contributions of individual nucleases to Okazaki fragment processing in the context of non-nucleosomal DNA. **A.** Distribution of Okazaki fragment 5’ termini around binding sites for Abf1, Reb1 and Rap1 (MacIsaac et al., 2006; Smith and Whitehouse, 2012). As in Figure 2, WT, single mutants, double mutants and the triple mutant are displayed in separate graphs for clarity. Data are presented such that Okazaki fragment synthesis proceeds from left to right. Data are normalized to the median signal in the range, and smoothed to 5 bp. **B.** Distribution of Okazaki fragment 3’ termini around binding sites for Abf1, Reb1 and Rap1, as in (A).

In both wild-type and single nuclease depletions, we observed a prominent peak of Okazaki fragment 5’ and 3’ termini immediately upstream of a meta-binding site for Abf1/Reb1/Rap1 (Fig. 3 A&B). However, the *RAD27-FRB exo1Δ* strain showed significantly reduced termination on the replication-fork proximal edge of the transcription-factor binding sites relative to the wild-type strain. The *RAD27-FRB DNA2-FRB; exo1Δ* strain generated slightly fewer fragments terminating at transcription-factor binding sites than the *RAD27-FRB exo1Δ* double mutant, suggesting that the decreased strand-displacement on non-nucleosomal DNA in the absence of both Rad27 and Exo1 can be further reduced by removal of Dna2. These data are consistent with a globally similar distribution of nuclease activity on non-nucleosomal DNA to that observed within nucleosomes – i.e. major redundant roles for Rad27 and Exo1 and a limited role for Dna2. However, the distribution of Okazaki fragment termini around TF binding sites in both *RAD27-FRB* and *exo1Δ* strains closely resembles that of a wild-type strain (Fig. 3A&B). Therefore, our data suggest more extensive redundancy of Rad27 and Exo1 outside nucleosomes than on nucleosomal templates.

### Uncoupling of Okazaki fragment processing from DNA synthesis

To investigate the extent to which the processing and ligation of the lagging strand can be uncoupled from its synthesis *in vivo*, we constructed strains in which expression of *RAD27*, *DNA2*, *EXO1*, or *CDC9* was driven by the *CLB2* promoter, and the protein fused to the N-terminal degron of the Clb2 protein. Expression of such *CLB2-*tagged constructs is limited to very late S- and G2 phase (Karras and Jentsch, 2010). Myc-tagged versions of Clb2-Cdc9, Clb2-Rad27, Clb2-Exo1 and Clb2-Dna2 were detectable by Western blot: all proteins displayed the anticipated cell cycle expression profile, with expression limited to late S/G2 (Fig. S2A-F). The lower band in the Cdc9 blot represents the mitochondrial isoform of Cdc9 (Willer et al., 1999), which does not cycle in a cell cycle-dependent manner.

Sporulation of a heterozygous diploid *CLB2-CDC9/CDC9* strain produced four viable spores per tetrad, indicating that expression of DNA ligase I exclusively at the end of S-phase and in G2 is sufficient to support viability (Fig 4A). Consistent with delayed ligation of Okazaki fragments when *CDC9* expression is restricted, the *CLB2-CDC9* strain transiently accumulated Okazaki fragments following release from G1 arrest which disappeared when ligation was enabled by *CLB2-CDC9* expression in late S/G2 (Fig. 4B). Sporulation of a *CLB2-RAD27/RAD27 CLB2-EXO1/EXO1* strain also generated four viable spores per tetrad, although *CLB2-RAD27 CLB2-EXO1* double mutant haploids grew slowly (Fig. 4C). *CLB2-RAD27* cells phenocopied the temperature-sensitivity and slow growth of *rad27Δ* cells, suggesting that these phenotypes arise due to a lack of Rad27 during S-phase (Fig. S2G). The observation that Rad27 and Exo1 together process the majority of Okazaki fragments in *S. cereivisiae* (Figs 2-3), but can maintain viability when expressed only after the bulk of replication has been completed, suggests that Okazaki fragment processing need not occur concurrently with replication.

**Figure 4.**
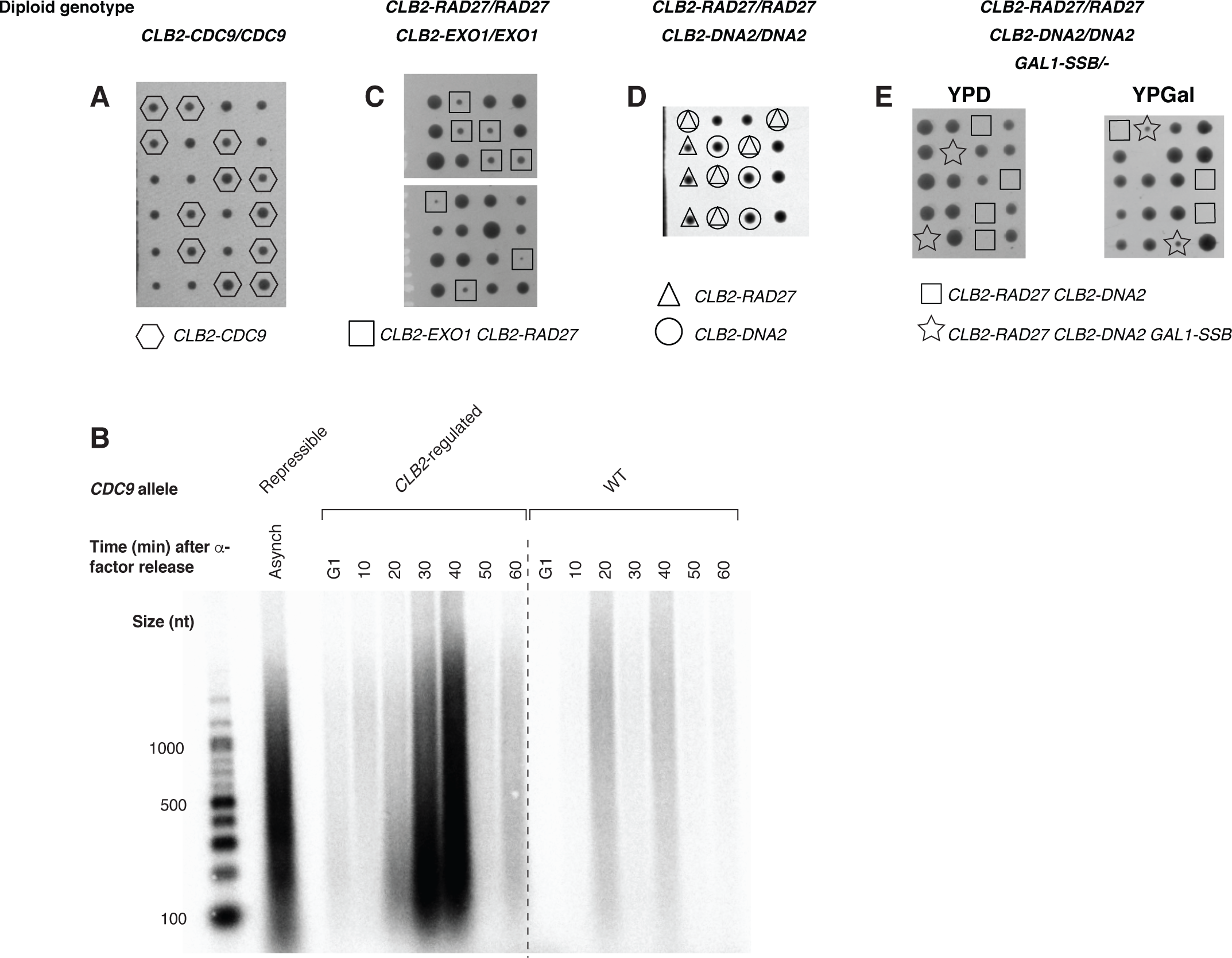
Uncoupling of Okazaki fragment processing and ligation from bulk DNA replication *in vivo*. **A.** Tetrads from sporulation of a diploid *CLB2-CDC9/CDC9* strain were dissected onto YPD and allowed to grow for 72h. **B.** *CLB2-CDC9* or wild-type cells were synchronized using alpha-factor, and released into fresh YPD for the indicated time. DNA was purified and end-labeled as in Fig. 1B. A sample from an asynchronous culture carrying a doxycycline repressible allele of *CDC9* is shown for comparison. Repression was for 2.5h as previously described (Smith and Whitehouse, 2012). **C&D.** Tetrads from sporulation of a diploid strain heterozygous for (C) *CLB2-RAD27* and *CLB2-EXO1* or (D) *CLB2-RAD27* and *CLB2-DNA2* were dissected onto YPD and allowed to grow for 72h. **E.** Tetrads from a diploid strain heterozygous for *CLB2-RAD27*, *CLB2-DNA2* and *GAL1-SSB* were dissected onto either YPD or YPGal plates, as indicated, and allowed to grow for 72h.

Sporulation of a *CLB2-RAD27/RAD27 CLB2-DNA2/DNA2* strain produced a mixture of viable and inviable spores (Fig. 4D). Colonies formed from *CLB2-DNA2* spores have an essentially wild-type growth phenotype (Fig. 4D). Thus, the essential contribution of Dna2 to viability can apparently be executed after the majority of the genome has been synthesized. In contrast to *CLB2-RAD27 CLB2-EXO1* double mutants, *CLB2-RAD27 CLB2-DNA2* haploids are inviable (Fig. 4D). Therefore, although Exo1 apparently contributes to the overall processing of Okazaki fragments to a greater extent than Dna2 (Fig. 2), at least some contribution from an endonucleolytic flap cleavage pathway (i.e. Dna2 or Rad27) is required for cells to progress through S-phase. We reasoned that the requirement for endonucleolytic Okazaki fragment flap processing during S-phase could arise from the accumulation of a small number of long, unprocessed flaps that might persist in *CLB2-RAD27 CLB2-DNA2* cells (Liu et al., 2017). These long flaps could deplete cellular RPA pools, leading to checkpoint-mediated cell cycle arrest (Byun et al., 2005) and/or catastrophic failure of replication (Toledo et al., 2013). To test whether excessive exposure of single-stranded DNA was the underlying cause of death in *CLB2-RAD27 CLB2-DNA2* cells, we expressed the *E. coli* single-stranded binding protein SSB from a galactose-inducible promoter. *CLB2-RAD27 CLB2-DNA2 GAL1-SSB* cells were inviable when grown on YPD to repress SSB expression, but viable when SSB was induced by growth on 2% galactose (Fig. 4E). Thus, expression of a generic single-strand binding protein restores viability when neither Rad27 nor Dna2 is expressed during early or mid S-phase, indicating that endonucleolytic processing of the lagging strand can be deferred until after the majority of replication has been completed. Because we observed that Okazaki fragment ligation can be similarly deferred (Fig 4A-B), we conclude that both steps of lagging-strand processing can be uncoupled from ongoing DNA replication in *S. cerevisiae*.

## DISCUSSION

In this study, we investigated Okazaki fragment processing in the absence of all combinations of the Okazaki fragment nucleases Rad27, Dna2 and Exo1 by carrying out a systematic *in vivo* analysis of Okazaki fragment size, ligatability and terminus location. Similarly to observations *in vitro* (Garg et al., 2004; Stodola and Burgers, 2016), we observe that predominantly ligatable Okazaki fragments are generated even in the absence of all three nucleases (Fig. 1), and that strand displacement synthesis by Pol δ is limited by the absence of nucleolytic Okazaki fragment cleavage (Figs. 2&3). However, a strain in which the expression of both flap endonucleases, Rad27 and Dna2, is restricted until after bulk DNA synthesis can only survive when a single-stranded binding protein is overexpressed. The large majority of Okazaki fragment termini apparently remain juxtaposed as ligatable nicks even without significant nuclease processing. However, the inviability of *CLB2-RAD27 CLB2-DNA2* cells suggests that some long flaps form in the absence of endonucleolytic Okazaki fragment processing, and that the formation of these structures is toxic via either RPA depletion, persistent checkpoint activation, or both.

Our data suggest that Rad27 and Exo1 have redundant activity in Okazaki fragment 5’ flap processing (Figs 2&3). If each nuclease acted on a preferred set of substrates, the most likely outcome would be a bimodal distribution of Okazaki fragment termini in single mutant strains. Substrates normally cleaved by the depleted/deleted nuclease would show a change in Okazaki fragment terminus location while those normally cleaved by the other nuclease would be unaffected. We do not observe such a bimodal distribution: instead, removal of either nuclease appears to globally impact lagging-strand processing and strand displacement by Pol δ. Thus, we propose that distinct Okazaki nucleases do not have rigid substrate preferences, but instead compete to cleave lagging-strand substrates *in vivo*. This model is consistent with the observed suppression of replication-associated *rad27Δ* phenotypes by overexpression of Exo1 (Tishkoff et al., 1997).

Although Rad27 and Exo1 can redundantly process lagging-strand 5’ flaps, removal of either nuclease leads to a global reduction in strand-displacement synthesis by Pol δ, especially in the context of nucleosomes (Figs. 2&3). The absence of Rad27 reduces strand displacement more than the absence of Exo1. Because each Okazaki fragment is initiated by the error-prone Pol α, such a widespread reduction in strand displacement would most likely increase the extent to which DNA synthesized by Pol α is transmitted to daughter cells. Both *rad27Δ* (Reagan et al.,1995) and *exo1Δ* (Tishkoff et al., 1997) strains are mutators, and the phenotype of *rad27Δ* is notably more severe than that of *exo1Δ* Given the correlation between the strength of the mutator phenotype and the extent to which strand-displacement by Pol δ is reduced, we speculate that retention of genomic DNA synthesized by Pol α underlies the elevated mutation rates observed when these genes are absent. Consistent with this model, temperature-sensitive *dna2* strains are not mutators (Budd et al., 2005) and depletion of Dna2 does not significantly alter global strand displacement *in vivo* (Figs. 2&3).

We observe that the contribution of Dna2 to Okazaki fragment processing is extremely limited in the uniquely mappable regions of the genome that can be assayed using our sequencing methodology (Figs. 2&3). In addition, *CLB2-DNA2* cells do not have a detectable growth defect (Fig. 4D). Therefore, although the nuclease activity of Dna2 is essential for viability (Budd et al., 2000), the absence of Dna2 during S-phase is not obviously detrimental to exponentially dividing *S. cerevisiae* cells. It is possible that Dna2 is required for the cleavage of only a small number of Okazaki fragment 5’ flaps – either long flaps generated by low levels of ongoing strand displacement without nuclease cleavage, or those in specific, late-replicating repeat regions such as telomeres (Budd and Campbell, 2013; Markiewicz-Potoczny et al., 2018) or the rDNA repeat (Villa et al., 2016).

## ACKNOWLEDGEMENTS

We thank the NYU Gencore for assistance with TapeStation and sequencing, and members of the Smith lab for helpful discussions. This work was supported by grants from the NIH (R01 GM114340), the March of Dimes (FY15-BOC-2141) and the Searle Scholars Program to D.J.S.J.S.O was supported by an American Cancer Society - New York Cancer Research Fund postdoctoral fellowship (PF-16-096-01-DMB).

## DATA AVAILABILITY

Sequencing data have been submitted to the GEO under accession number GSE118078

## AUTHOR CONTRIBUTIONS

M.K. and J.S.O. performed research. R.Y. and J.S.O. analyzed sequencing data. D.J.S. supervised the study.

## METHODS

### Yeast strains, cell growth and spot tests

Yeast strains were all W303 RAD5+. The wild type strain genotype is *matA, tor1-1::HIS3, fpr1::NatMX4, RPL13A-2xFKBP12::TRP1, CDC9-FRB::KanMX6*. Additional FRB tagging, gene deletions, Clb2 promoter insertion and myc tagging were performed by PCR-mediated replacement, and introduced into the parent strain by crossing. At least two independent replicates of each strain were selected after tetrad dissection.

For *GAL1-SSB* strains, the pRS405 integrating vector containing a codon-optimized open reading frame encoding *E. coli* SSB under the control of the *GAL1* promoter was integrated at the *LEU2* locus. Stains were grown at 30ºC in YPD unless otherwise specified. Rapamycin was added to a final concentration of 1 μg/ml in liquid media or 2 μg/ml in solid media. For spot tests, cells were washed and diluted to an OD of 1. Cells were plated at a 1:10 dilution series and grown overnight at 30 or 37˚C as specified.

### Cell-cycle synchronization and western blotting

200ml of exponential phase cell cultures at OD 0.3-0.4 was synchronized in G1 using 5μg/ml alpha factor. Cells were released into S-phase and harvested at different time points. Growth was immediately stopped by addition of ice cold water and cells were centrifuged at 4ºC. Cells were resuspended for 5 minutes in 2M lithium acetate at 4ºC, pelleted, resuspended for 5 minutes in 400mM sodium acetate at 4ºC, pelleted again and finally resuspended in 1X Laemmli buffer plus 5% beta-mercaptoethanol. After 5min of boiling at 100ºC, the lysates were centrifuged for 5 minutes at top speed and transferred to new tubes prior to loading on a 10% SDS-Page gel. After migration at 100V, samples were transferred to a PVDF membrane, blocked with 5%milk in TBS-0,1% Tween and probed with a C-Myc antibody (Genscript A00173-100). Loading controls were done by Coomassie staining of gels loaded with identical amounts of sample and run alongside gels for Western blots.

### Fluorescence-activated cell sorter (FACS) analysis

150 *μ*l of cells were harvested during the S-phase release at different timepoints and 350*μ*l of 100% ethanol was added to fix the cells overnight at 4ºC. Cells were pelleted, resuspended in 50mM sodium citrate plus Rnase A and incubated at 50ºC for an hour. After addition of proteinase K and another hour of incubation, cells were labeled with SYTOX green and stored at 4ºC before analysis using a Becton Dickinson Accuri.

### Okazaki fragment analysis

Okazaki fragments were purified and end-labeled essentially as described (Smith and Whitehouse, 2012). Briefly, DNA concentration was normalized by Qubit. 650 ng of DNA per lane was treated with Klenow (exo-) (NEB) and α-^32^P dCTP for 30 min at 37˚C in NEBuffer 2. Unlabeled nucleotides were removed using Illustra G50 columns (GE Healthcare) and samples were separated for 5h at 77V in 1.3% alkaline agarose gels. DNA was transferred to a Nylon membrane overnight and visualized using a phosphorimager.

For *in vitro* ligation and Taq cleavage experiments, samples were incubated in 1X DNA ligase buffer (NEB). 1μl of Taq DNA polymerase (NEB) was added to indicated samples, and incubated for 15min at 50ºC. Subsequently, 2μl of T4 DNA ligase (NEB) was added and samples were incubated for 90 min at room temperature before phenol extraction and end-labeling.

Okazaki fragment purification and sequencing was carried out as previously described (Smith and Whitehouse, 2012). Paired-end sequencing (2 × 75 bp) was carried out on an Illumina Next-seq 500 platform.

---

**Figure S1.**
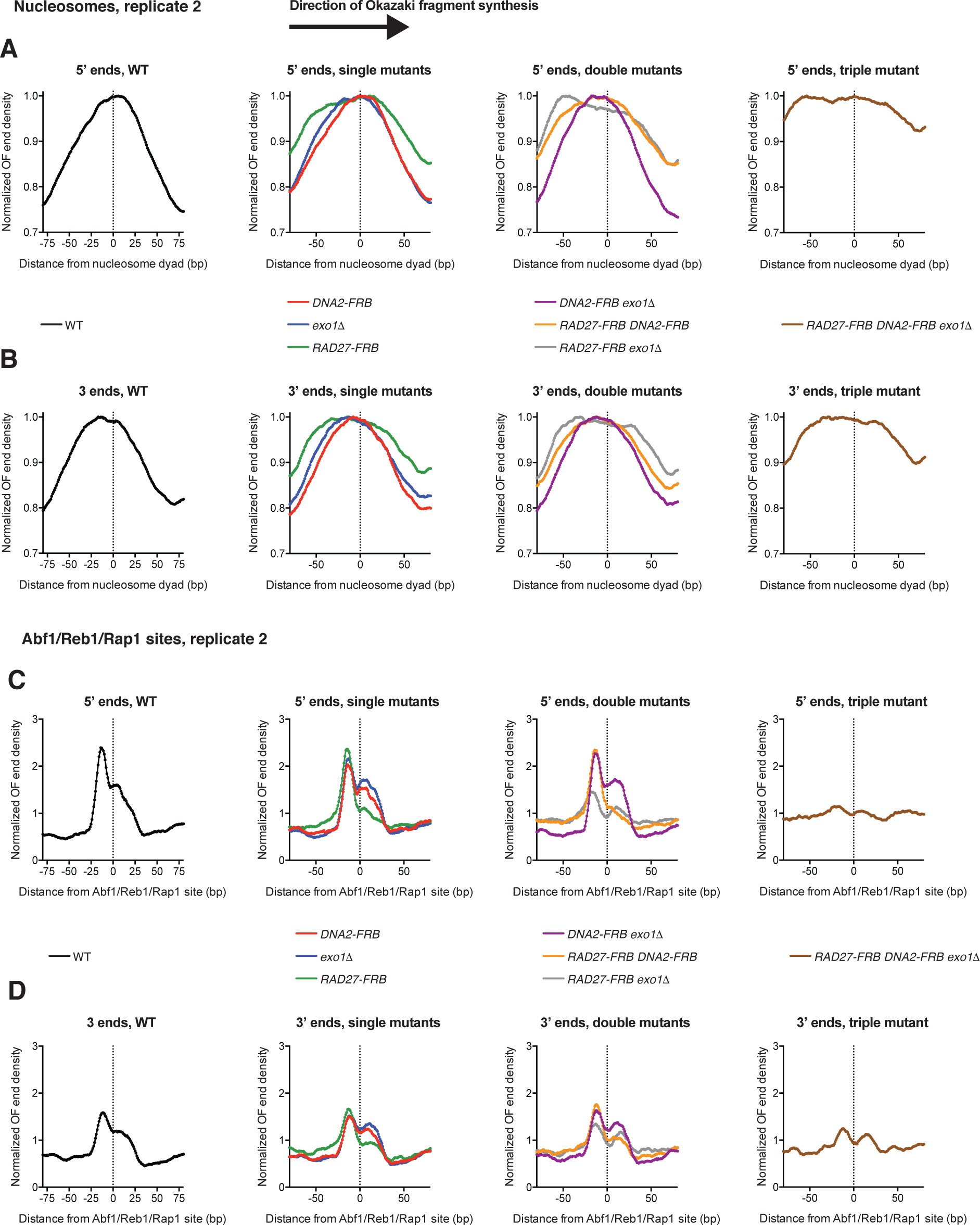
Sequencing data replicate comparisons. **A.** Distribution of Okazaki fragment 5’ termini around nucleosome dyads, as in Fig. 2A, for a second biological replicate of each strain. **B.** Distribution of Okazaki fragment 3’ termini around nucleosome dyads, as in Fig. 2B, for a second biological replicate of each strain. **C.** Distribution of Okazaki fragment 5’ termini around Abf1/Reb1/Rap1 sites, as in Fig. 3A, for a second biological replicate of each strain. **D.** Distribution of Okazaki fragment 3’ termini around Abf1/Reb1/Rap1 sites, as in Fig. 3B, for a second biological replicate of each strain.

**Figure S2.**
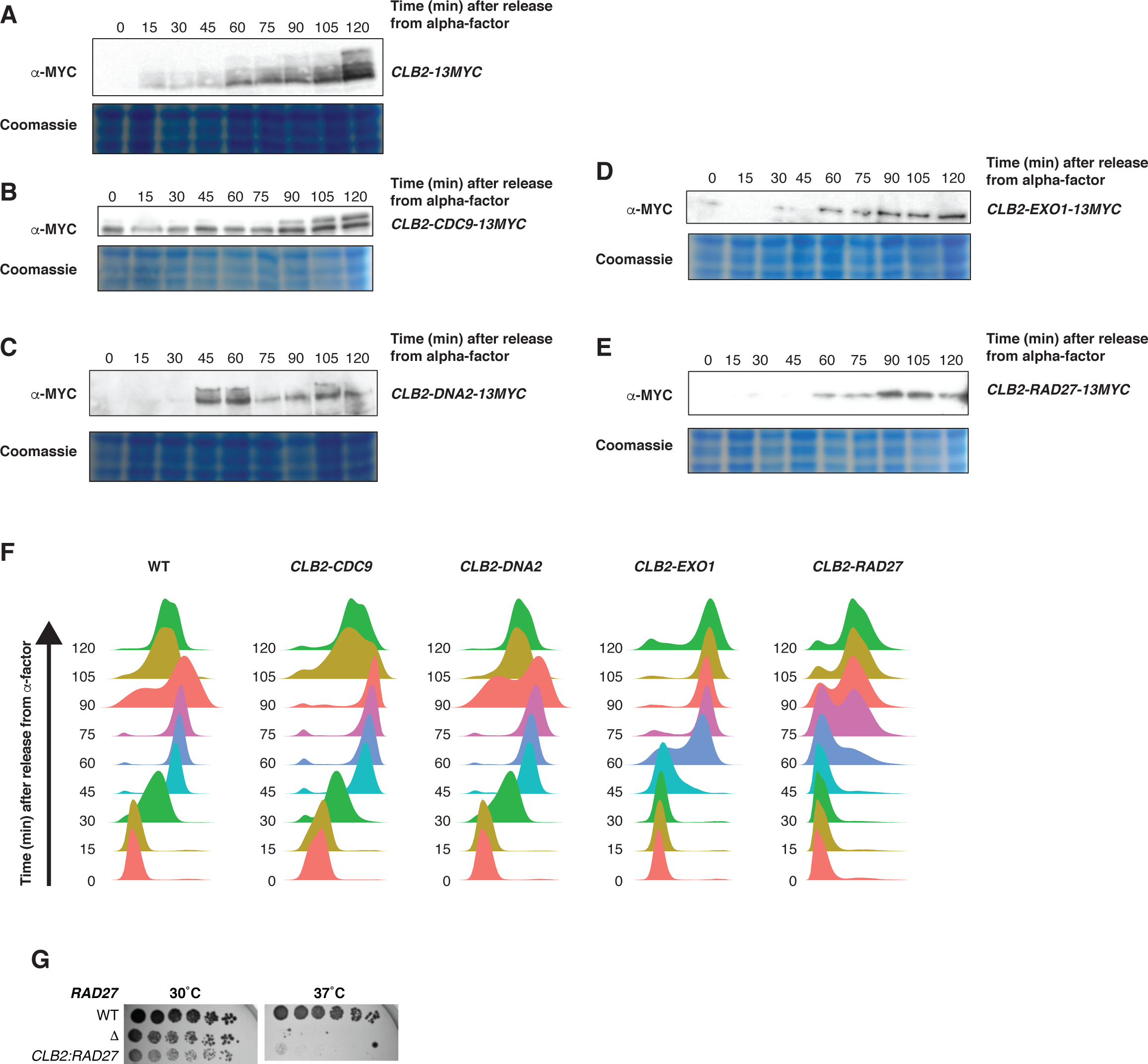
Expression and cell-cycle analysis for strains expressing Clb2-regulated Okazaki-fragment processing enzymes. **A-E.** Anti-Myc Western blots from strains released from alpha-factor arrest for the indicated time. *CLB2* (A), *CLB2-CDC9* (B), *CLB2-DNA2* (C), *CLB2-EXO1* (D) or *CLB2-RAD27* (E) were C-terminally tagged with 13xMyc. Identically loaded Coomassie-stained gels are shown as loading controls. The lower band in (B) represents the mitochondrial isoform of Cdc9, which uses a distinct translational start site and does not cycle. **F.** DNA content, assayed by flow cytometry, for samples analyzed in Fig. S2A-E **G.** Spot tests on YPD at 30˚C or 37˚C, as indicated, comparing the growth of WT, rad27Δ and *CLB2-RAD27* cells.

